# Plant production of high affinity nanobodies that block SARS-CoV-2 spike protein binding with its receptor, human angiotensin converting enzyme

**DOI:** 10.1101/2022.09.03.506425

**Authors:** Marco Pitino, Laura A. Fleites, Lauren Shrum, Michelle Heck, Robert G. Shatters

## Abstract

Nanobodies® (V_HH_ antibodies), are small peptides that represent the antigen binding domain, V_HH_ of unique single domain antibodies (heavy chain only antibodies, HcAb) derived from camelids. Here, we demonstrate production of V_HH_ nanobodies against the SARS-CoV-2 spike proteins in the solanaceous plant *Nicotiana benthamiana* through transient expression and their subsequent detection verified through western blot. We demonstrate that these nanobodies competitively inhibit binding between the SARS-CoV-2 spike protein receptor binding domain and its human receptor protein, angiotensin converting enzyme 2 (ACE2). We present plant production of nanobodies as an economical and scalable alternative to rapidly respond to therapeutic needs for emerging pathogens in human medicine and agriculture.

## 0 Introduction

Severe acute respiratory syndrome coronavirus 2 (SARS-CoV-2) is a member of the subfamily *Coronaviridae* in the family *Coronaviridae* and the order *Nidovirales*. Pathogenic viruses in this subfamily cause severe respiratory syndrome in humans. SARS-CoV-2 is related to SARS-CoV-1 and Middle Eastern Respiratory Syndrome (MERS-CoV), which emerged in humans in 2003 and 2012, respectively. SARS-CoV-2 is responsible for the 2019 pandemic and COVID-19 disease (Huang, Wang et al. 2020). COVID-19 disease results in a range of outcomes, ranging from asymptomatic infection to patient death. To date, global vaccinations for SARS-CoV-2 protections are underway, but additional treatments are needed to prevent infection among naïve and even vaccinated individuals. Tiered prevention efforts have been shown to reduce transmission and severity of disease outcome.

Coronaviruses are positive-sense, single-stranded RNA viruses with spherical virions bound by a membrane envelope that are 100-160nm in diameter. The 3’ end of the viral genome encodes the structural proteins, including the envelop glycoprotein spike (S), envelop (E), membrane (M) and nucleocapsid (N). Inserted into the membrane envelop are ~25 copies of the homotrimeric transmembrane spike glycoprotein (spike protein) as a clover-shaped trimer, with three S1 heads and a trimeric S2 stalk (Benton, Wrobel et al. 2020). The receptor binding domain (RBD) is situated atop each S1 head (Nishima and Kulik 2021). The RBD is responsible for entry into host cells (Wang, Zhang et al. 2020, Jackson, Farzan et al. 2022) via interaction with the protein angiotensin converting enzyme 2 (ACE2), the interaction which also determines the viral host range (Yan, Zhang et al. 2020). Studies have shown a higher affinity for SARS-CoV-2 to ACE2 as compared to ACE1, further supporting its role in transmission and virulence (Samavati and Uhal 2020). Highly transmissible viral variants, such as Delta and Omicron variant, have been selected for during the pandemic and exhibit mutations in the RBD (Li, Lai et al. 2021) (Saxena, Kumar et al. 2022). Thus, interactions between ACE2 and the RBD are attractive targets for the development of novel anti-viral therapies.

Nanobodies represent a promising new therapy for the treatment of viral diseases, including COVID-19. A pubmed search for SARS-CoV-2 and nanobody brings up a total of 21 peer-reviewed publications (Esparza, Martin et al. 2020). Nanobodies, also referred to as V_HH_, are produced by animals in the camelid family, which include llamas and alpacas. Coined by the popular press as mini-antibodies (Deyev and Lebedenko 2009), these IgGs are less than 15 kDa and are comprised of an unpaired heavy-chain variable domain. Nanobodies have been reported to bind antigens with affinities equivalent to a conventional IgG (Gonzalez-Sapienza, Rossotti et al. 2017, Asaadi, Jouneghani et al. 2021). Nanobodies are also under development for the control of at least two crop diseases: grapevine fanleaf virus in cultivated wine grapes (Yan, Zhang et al. 2020), botrytis, and detection against a range of other plant pathogens (Njeru and Kusolwa 2021).

These antigen-binding proteins, derived from single-chain camelid antibodies, are significantly smaller in size compared to conventional antibodies with a molecular mass of 12-15 KDa (conventional antibodies are ~150 kDa). Key features of nanobodies that make them attractive alternatives to conventional antibodies include their high affinity, specificity, solubility, thermostability and mobility. Production of nanobodies is typically done by expression of the gene in *E.coli*; however, a potentially more effective method is currently being studied based on plant expression.

We represent a team of agricultural scientists developing sustainable and biologically-based solutions to pathogens of economic importance in crop production. As part of this research, we developed a low-cost, plant-based method of producing proteins that could be used to solve agricultural pathogen problems in agricultural production settings. As a proof-of-concept, we describe the production of a RBD nanobody in a plant expression system. The benefits of producing therapeutics in plants justify considering plants to mass produce COVID-19 protein-based therapies.

## 1 Materials and Methods

### 1.1 Construct Design

A total of four constructs were designed for experimentation with plant expression of COVID-19 nanobodies. The methionine start codon of the SARS-CoV2 nanobody protein sequence (NIH-CoVnb-112; Esparza et al. 2020) was removed and replaced with an N-terminal signal peptide sequence for protein secretion into the apoplast and a 6x histidine tag at the C-terminus (SP-CoV19_his). An analogous negative mutated control construct was also designed, such that the amino acids spanning the three complementarity determining regions (CDR1, CDR2, CDR3) of the native SARS-CoV2 nanobody sequence were scrambled using a random number generator SP-*mCov19*_his; (Fig. S1). Disruption of the CDR regions was expected to abolish the interaction with the receptor binding domain of the viral spike protein.

Two more variants of the SARS-CoV2 nanobody construct were made as fusions to monomeric enhanced green fluorescent protein coding sequence (EGFP): one with an N-terminal 6× histidine tag (SP-his_CoV19-GFP), and a second with a C-terminal 6× histidine tag (SP-CoV19_his-GFP; (Fig. S1). This module was followed in both constructs by EGFP, with a P2A site (ribosome skipping sequence allowing both CoV19-variants and EGFP to be produced as separate proteins) inserted between the nanobody module and EGFP sequence. All constructs were codon-optimized for expression in the Solanaceae using an online tool provided by Integrated DNA Technologies (IDT, Illinois, USA; https://www.idtdna.com/CodonOpt) prior to uploading to the online ordering portal for Codex DNA (La Jolla, CA). A 40bp span of nucleotide sequence homologous to the recipient vector pNANO was added to the 5’ and 3’ ends of the constructs to enable cloning with the BioXP 3250 system (Codex DNA, La Jolla, CA).

### 1.2 Construct Generation and Bacterial Transformation

The plasmid backbone was linearized by sequential digestion with *Sma*I and *Spe*I (New England Biolabs, Ipswich, MA, USA), and gel purified from 0.8% SeaPlaque GTG Agarose (Lonza, Rockland, ME) using a phenol:chloroform:isoamyl alcohol extraction method followed by overnight precipitation at −20°C in 100% ethanol, 0.3M NaOAc, pH 5.0. The purified, precipitated DNA was washed with 70% ethanol, dried briefly, and resuspended in sterile nuclease free water. The final constructs were generated in an overnight run on the BioXP 3250 system (Fig. 1A) (Codex DNA, La Jolla, CA), an automated synthetic biology platform for DNA fragment assembly and cloning.

**Figure 1.**
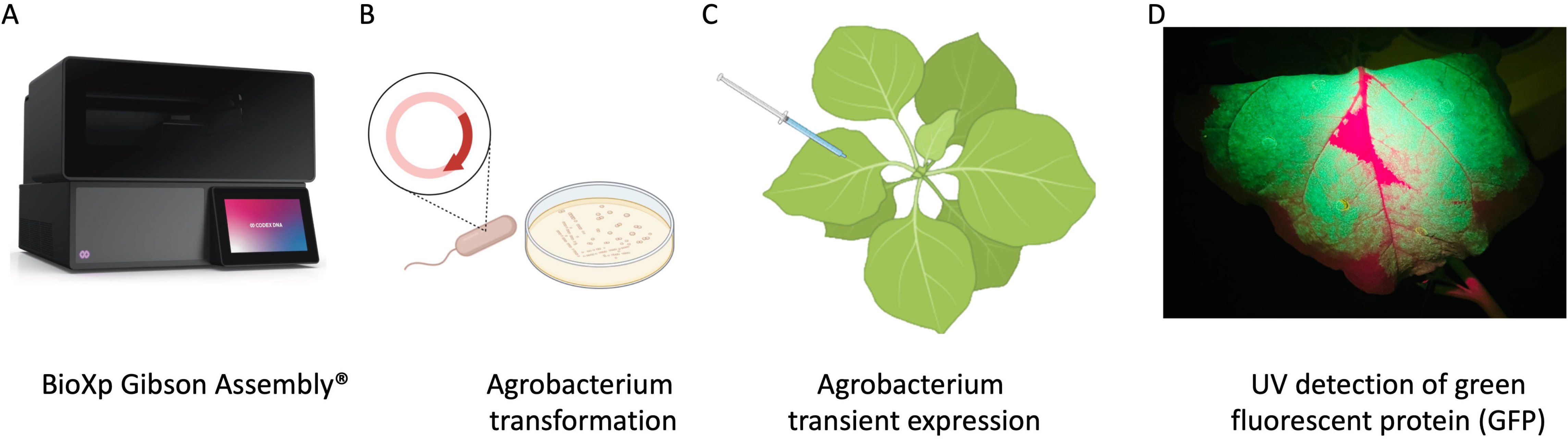
Schematic representation of workflow of production of nanobodies in plant system. A) Cloning with the BioXP 3250 Gibson Assembly®. B) Agrobacterium transformation. C) Agrobacterium infiltration was performed by using 1 mL needleless syringes to inject bacteria into the abaxial side of the leaves at OD_600_ 0.3. D) P2A sequence was used for generating multiple separate proteins from a single mRNA, GFP included in the sequence allowed prescreening of high expression protein in leaves using UV light.

*Agrobacterium tumefaciens* EHA105 was electroporated with the BioXP products and grown on LB supplemented with kanamycin (100μg/ml) for three days (Fig. 1B). Colonies were screened using colony PCR and sequence verified prior to transient expression and purification in *N. benthamiana*.

### 1.3 Plant Growth and Agroinfiltration

*N. benthamiana* plants were grown under the greenhouse conditions and used at 4-5 weeks old for transient expression using plant infiltration with *Agrobacterium* EHA105, which mechanically delivers the bacteria to the plant’s extracellular matrix (apoplast) (Kapila, De Rycke et al. 1997) (Fig.1C). *Agrobacterium* EHA105 harboring pNANO plasmid was cultured overnight in 5 mL of LB media with 100μg ml^-1^ of kanamycin. Overnight culture was pelleted and resuspended in infiltration buffer (10mM MgCl2, 10mM MES, 400 μM acetosyringone) at optical density at 600 nm (OD_600_) 0.3. For each construct, leaves were infiltrated with the bacterial suspension and set in greenhouse for duration of experiment (Fig. 1C). Two days post infiltration (2 dpi), leaves were manually excised from the plants using a sterile blade and processed for total protein extraction and purification.

### 1.4 Protein Extraction and Purification

*N. benthamiana* leaves were observed under UV light for GFP expression and harvested at 2 dpi (days post infiltration) (Fig. 1D) followed by homogenization in liquid nitrogen. Total plant proteins were extracted using native buffer (10mM Tris/HCl pH 7.5, 150mM NaCl, 0.5mM EDTA, 1% [v/v] P9599 Protease Inhibitor Cocktail [Sigma-Aldrich], 1% [v/v] IGEPAL CA-630 [Sigma-Aldrich]). A total of 5mL of extraction buffer per gram of leaf tissue was used. Samples were clarified by centrifugation at 4°C at 3000 rcf. The supernatant was filtered through a 40 μm nylon cell strainer (Becton Dickinson Labware, Franklin Lakes, NJ, US) and then used for purification process utilizing Ni-NTA agarose columns (Thermo Scientific, Rockford, US), following manufacturers guidelines. Briefly, imidazole binding buffer (20mM sodium phosphate, 10mM imidazole, 0.5mM NaCl, pH 7.4) was used to equilibrate, bind, and wash the columns. To elute product of interest, (20mM sodium phosphate, 500mM imidazole, 0.5mM NaCl, pH 7.4) was used.

### 1.5 SDS-PAGE and Western Blotting

Samples were denatured and reduced using 5× Lane Marker Reducing Sample Buffer (Thermo Scientific, Rockford, IL, US), boiled at 95°C for 10 minutes, then stored on ice. Gradient 4-20% precast polyacrylamide gels (Bio-Rad Laboratories, Hercules, CA) were loaded into electrophoresis tank (Bio-Rad Laboratories) and filled with 1× Tris/Glycine/SDS buffer. Kaleidoscope ladder (Bio-Rad Laboratories) was loaded into the first well (5uL), and each sample was loaded into every other well. 25μL per sample were used for Coomassie staining, and 10uL per sample were used for immunoblotting. Electrophoresis was run following manufacturing guidelines with a powerpack (Bio-Rad Laboratories). One gel was stained with Coomassie blue, while the other gel was transferred to a nitrocellulose membrane using the Trans-Blot Turbo Transfer system following manufacturer guidelines (Bio-Rad Laboratories). The nitrocellulose membrane was removed and placed in 1X Casein blocker for one hour on rotator followed by incubation with a 1:1000 dilution of his HRP-conjugated antibodies (Proteintech, Rosemount, IL, US) for 1 hour room temperature. The membrane was washed in 1X TBS three times in 10-minute intervals. ECL substrate (Bio-rad Laboratories) that consists of 1 mL peroxide and 1mL luminol enhancer were spread onto membrane and left for 5 minutes before observation using ChemiDoc imager (Bio-rad Laboratories).

### 1.6 Competitive ELISA binding screen for ACE2 and RBD

To verify the activity of recombinant nanobodies generated in plants, we conducted a competitive binding assay that measures inhibition of the interaction between the receptor binding domain (RBD) of the SARS-CoV-2 spike protein with the ACE2 receptor in the presence of the purified nanobodies. Purified nanobodies were diluted at 1μg/mL and 0.1 μg/mL concentrations in association with RBD proteins then added to the ACE2 coated plate (RayBiotech, Peachtree Corners, GA, US). Nanobodies and RBD proteins were incubated at room temperature for one hour to allow interaction. The assay plate was washed four times with a wash solution provided by ELISA kit (RayBiotech). HRP-conjugated Anti-IgG was added to plate post wash and incubated at room temperature for one hour with gentle shaking. After four additional washes, the plate was developed by addition of tetramethylbenzidine and stopped after 30 minutes of gentle shaking in the dark with stop solution (RayBiotech). Absorbency was measured immediately after adding stop solution at 450 nm on a plate reader (Citation 1 imaging reader, BioTek, Winooski, VT, US).

## 2 Results

### 2.1 Expression and purification of nanobodies for SARS-Cov-2 RBD in *N. benthamiana*

An initial test performed using SP-CoV19 his purified from transient expressing leaves showed an expected ~15KDa band confirming expression and purification. This band was visualized by Coomassie blue staining of an SDS-Page gel and western blot/immunodetection specific for the his-tag on the SP-CoV19 protein (Fig. S2). Next, we tested SP-his CoV19-GFP, SP-CoV19 his-GFP, SP-*mCov19* his sequences. GFP visualization of infiltrated leaves showed high levels of expression two days post infiltration and leaves were harvested at this time and used for purification. Bands on the Coomassie gel and western blot were visualized migrating between the 15 and 20 kDa marker bands corresponding to the size of the SP-his_CoV19-GFP, SP-CoV19_his-GFP and SP-*mCov19*_his sequences (Fig 2A,B).

**Figure 2.**
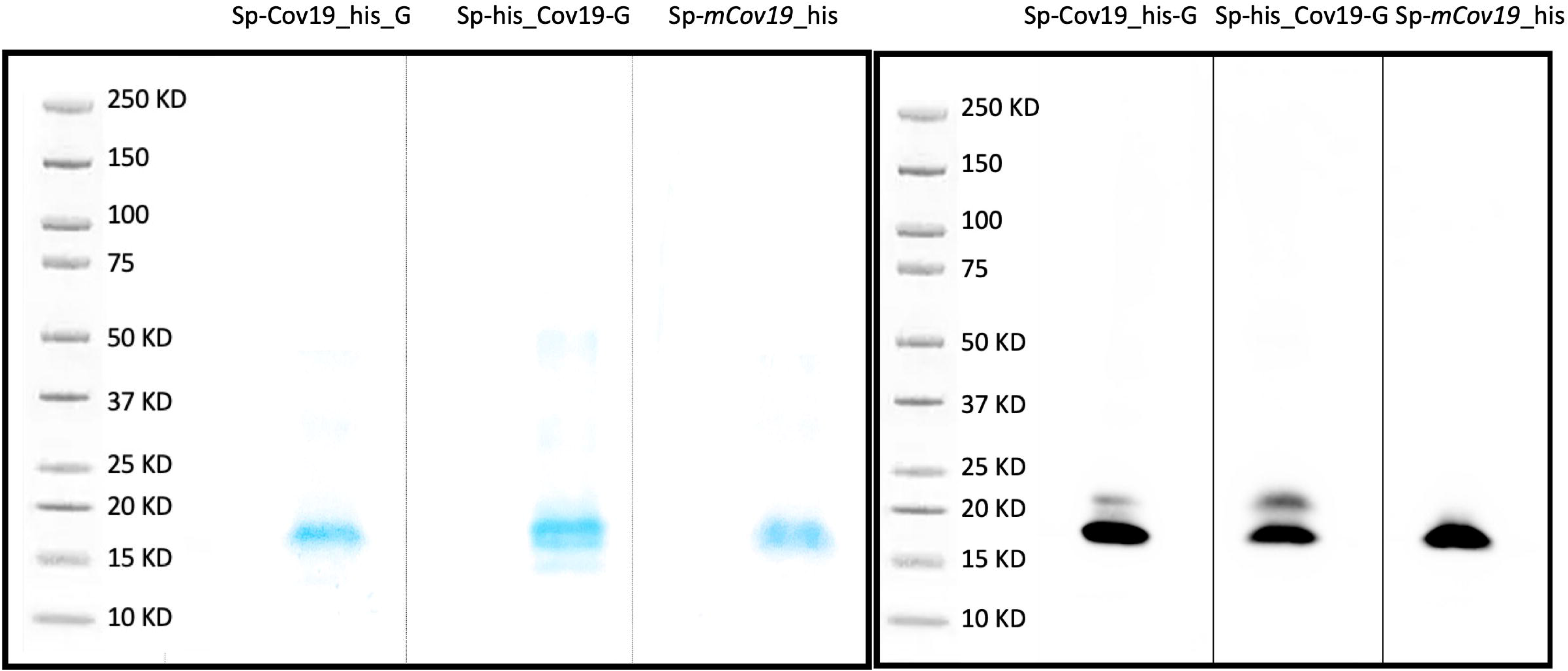
SDS PAGE and Western blot. A) Coomassie blue stain was used to verify purity of concentrated proteins and band size. **B)** Western blotting was carried out to detect the target purified SP-CoV19_his-GFP, SP-his_CoV19-GFP and negative control SP-*mCov19*_his using his antibodies.

### 2.2 Biological activity of SARS-Cov-2 nanobody with ACE2 competition assay

Next, we assessed the ability for plant produced nanobody to block ACE2 binding cells RBD interaction. To evaluate relative inhibition of RBD protein from binding to ACE2 a competitive ELISA inhibition assay was performed. RBD protein binding ACE2 was indicated by high colorimetric absorbance. Initial screening was performed using SP-CoV19 his at 100,10,1 and 0.1 μg/mL concentrations (Fig. S2C). Competitive ELISA assay indicated that 1μg/mL SP-CoV19 his inhibited interaction between ACE2 and RBD and was used for subsequent experiments. The same results were obtained using both SP-his CoV19-GFP and SP-CoV19 his-GFP, showing 100% inhibition between ACE2 and RBD at 1μg/ml. Inhibition was also observed at 0.1μg/ml with 60-70% inhibition (Fig. 3). In contrast, the mutated sequence SP-*mCov19*_his inhibited less than <20% at 1.0 μg/ml and 0% at 0.1 μg/ml (Fig. 3). These results showed that plant-produced SP-CoV19 his, SP-his_CoV19-GFP and SP-CoV19 his-GFP, but not SP-*mCov19* his, inhibit 100% ACE2 and RBD interactions at 1μg/mL similarly to previous published data with NIH-CoVnb-112 production in yeast system (Esparza et al. 2020).

**Figure 3.**
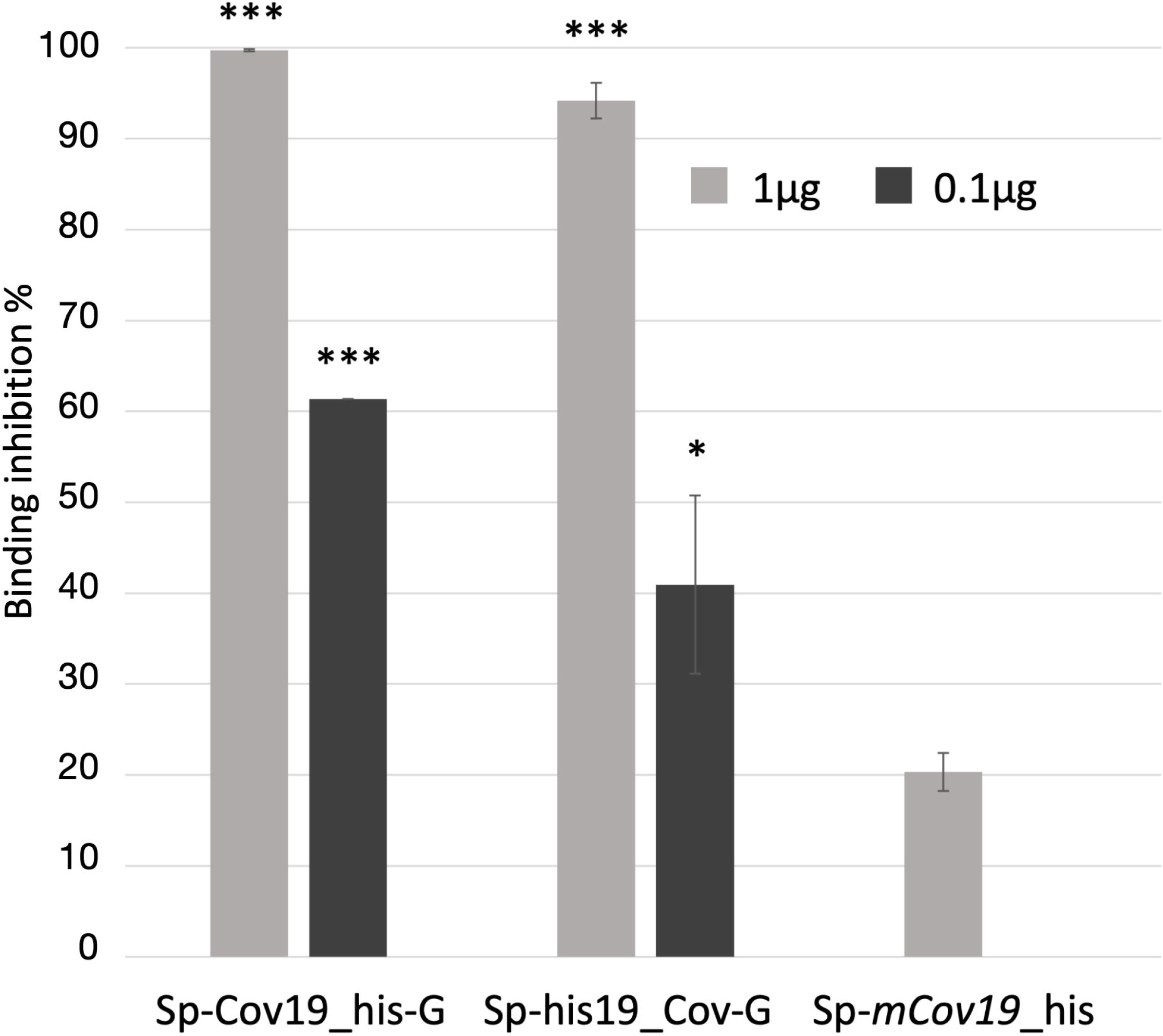
Competitive ELISA inhibition of ACE2 and RBD binding using anti RBD nanobodies. Competition binding assays were used to investigate whether the SP-Cov19_his-GFP and SP-his_CoV19-GFP blocked the binding of RBD to ACE2 compared to the mutant version SP-*mCov19*_his. SP-CoV19_his-GFP, SP-his_CoV19-GFP and SP-*mCov19*_his at 1μg/mL and 0.1μg/mL were incubated with RBD proteins, both SP-CoV19_his-GFP, SP-his_CoV19-GFP inhibited RBD bound to ACE-2 but SP-*mCoV19*_his at 1μg/mL. (Bars represent standard deviation, the asterisks indicate significant differences between conditions, * = *P* < .05, ** = *P* < .005, *** = *P* < .001).

## 3 Discussion

In this study, we provide proof of concept for in plant production of nanobodies that neutralize the interaction between the human ACE2 receptor and the SARS-CoV-2 spike protein RBD, a key step of the infection initiation process (Esparza, Martin et al. 2020). Binding inhibition was slightly reduced with a 10-fold dilution of the nanobody, consistent with previous reports for the same nanobody expressed in yeast (Esparza, Martin et al. 2020). Moreover, a modified nanobody with a scrambled RBD binding domain failed to inhibit binding at the lower concentrations used, demonstrating the binding specificity of the interaction between the plant-produced RBD-binding nanobodies and the RBD. The plant expression constructs used two features to aid in nanobody production: the use of a signal peptide targeting the nanobody to the plant apoplastic space and a self-cleaving P2A peptide. A signal peptide was added for future nanobody production in plant cell tissue culture systems, to support secretion of the nanobody through the cellular secretory pathway. The self-cleaving P2A peptide enabled production of functional nanobodies with concurrent fluorescent protein signal to monitor transient transformation events in *N. benthamiana* and to easily localize regions of nanobody production *in planta*. Plant tissue was harvested at 2 days post-infiltration, and thus not optimized for nanobody yield in this study. The His tag facilitated enrichment from the plant proteome but would need to be removed prior to the development of plant-based nanobody therapies for the treatment of human or other animal diseases. A recent example of production of RBD in planta exhibits suitable biochemical and antigenic features for use in a subunit vaccine platform (Demone, Nourimand et al. 2021). We posit that molecular farming of nanobodies, and other biologicals is an under-developed area for cost savings and increased global access for the production of protein and small molecule therapies.

Research using nanobodies in plants has been increasing rapidly in recent years (Dhama, Natesan et al. 2020, Wang, Yuan et al. 2021), including in the development of therapies and diagnostic tools for plant diseases. Plants offer several advantages for nanobodies expression over conventional expression platforms including their easy transformation, low risk of pathogen contamination and low cost for upscaling. In addition to injectable vaccines, new strategies are emerging and being developed to increase protection against COVID-19, for example a nasal spray-delivered nanobody offers a complementary barrier method to prevent virus acquisition into human epithelial cells in the airway. Nanobodies are 12–15 kDa single-domain antibody fragments that can be delivered by nebulizers and relatively easy and inexpensive to produce compared to other systems (Esparza et al. 2020). Previous cryo-electron microscopy studies showed SARS-CoV-2 spike protein and its interaction with the cell receptor ACE2, such binding triggers a cascade of events that leads to the fusion between cell and viral membranes for cell entry (Kirchdoerfer et al. 2018; Yuan et al. 2017; Gui et al. 2017). Because SARS-CoV-2 binding spike protein RBD and the host ACE2 receptor determines host susceptibility to the virus, interfering with that interaction might constitute a treatment option (Walls et al. 2020). Esparza and colleagues (2020) showed that NIH-CoVnb-112 candidate nanobodies blocked interaction between ACE2 and RBD “wild type” and 3 variant forms, also they showed that it retained structural integrity and potency after nebulization. A subset of these nanobodies fold in planta and retain the structural features necessary to interfere with protein interaction between ACE2 and the SARS-CoV-2 spike protein RBD. We demonstrate that nanobodies produced in plants retain proper folding and functionality comparably to a yeast production system supporting the use of plants as cost-effective production platforms for therapeutic needs with emerging pathogens, such as the SARS-CoV-2 virus.

## 5 Author Contributions

MH, MP and RS conceived of the idea and provided grant funding. LF developed and constructed the nanobody constructs. MP, LF and LS conducted the experiments. All authors contributed to data analysis and writing and editing the paper.

## 7 Acknowledgements

We would like to thank Carrie Vanderspool and Mant Acon for the plants used in this study. Mention of trade names or commercial products in this article is solely for the purpose of providing specific information and does not imply recommendation or endorsement by the U.S. Department of Agriculture.

## 8 Supplementary Material

Supplementary Material should be uploaded separately on submission, if there are Supplementary Figures, please include the caption in the same file as the figure. Supplementary Material templates

## Supplementary data

**Figure S1.** Multiple sequence alignment of protein coding sequences showing the overall structure of the various nanobody constructs. The complementarity determining regions (CDR) 1, 2, and 3, which were mutated in the negative control construct SP-*mCov19*_his, are annotated along with other major features.

**Figure S2. A)** Western blot serial elution: total of 5ml elution buffer with 500mM imidazole was used to recover of 1mL each sample B) Coomassie blue staining was used to validate purification after Ni-NTA using total protein and purified protein. **C)** Purified SP-CoV19_his samples were pulled together and concentrated using 10Kda size exclusion column to 500μl and tested with competitive ELISA at 100,10,1,0.1 μg/mL.

