## Supplementary figures and images for "Plant production of high affinity nanobodies that block SARS-CoV-2 spike protein binding with its receptor, human angiotensin converting enzyme"

### Fig. S1

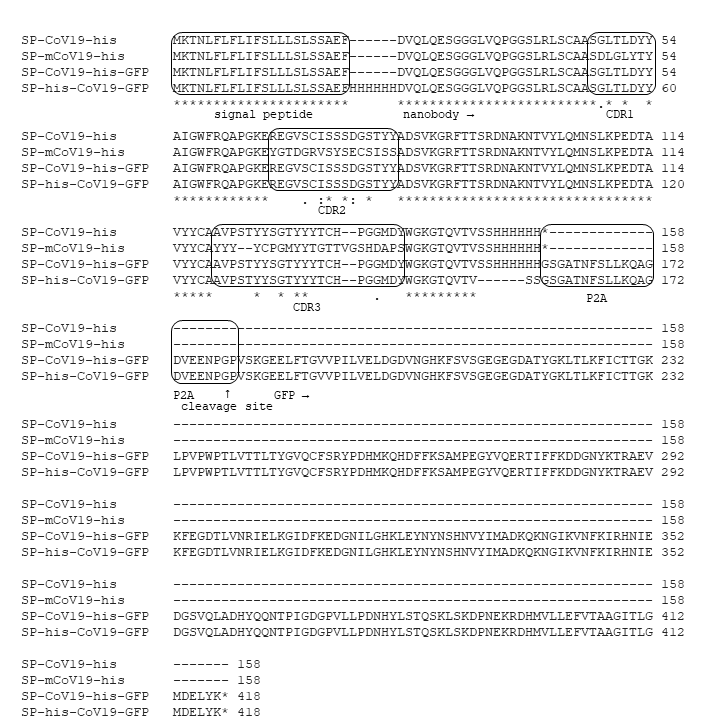

### Fig. S2

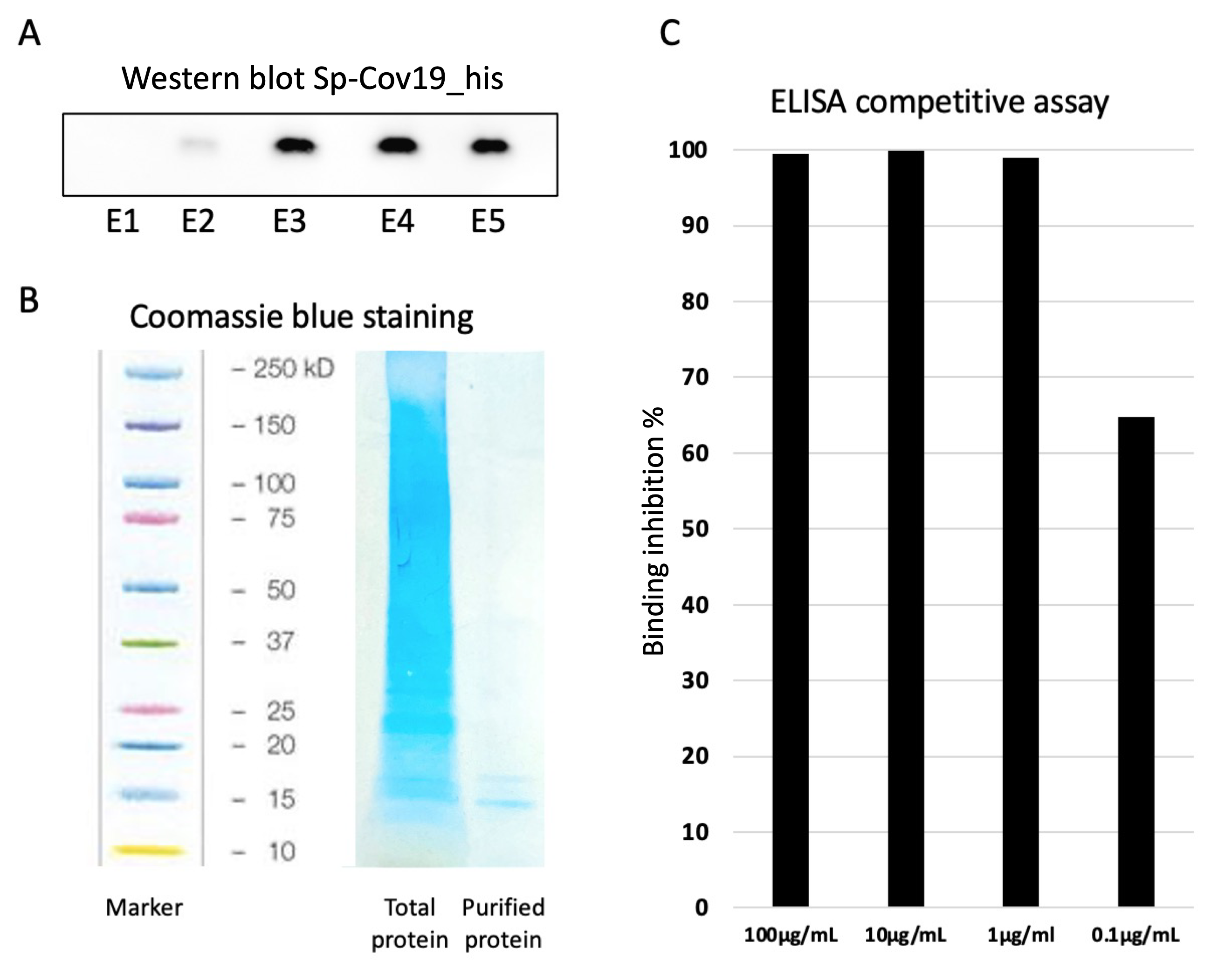
